# Physiology-informed regularization enables training of universal differential equation systems for biological applications

**DOI:** 10.1101/2024.05.28.596164

**Authors:** Max de Rooij, Balázs Erdős, Natal van Riel, Shauna O’Donovan

## Abstract

Systems biology tackles the challenge of understanding the high complexity in the internal regulation of homeostasis in the human body through mathematical modelling. These models can aid in the discovery of disease mechanisms and potential drug targets. However, on one hand the development and validation of knowledge-based mechanistic models is time-consuming and does not scale well with increasing features in medical data. On the other hand, more data-driven approaches such as machine learning models require large volumes of data to produce generalizable models. The integration of neural networks and mechanistic models, forming universal differential equation (UDE) models, enables the automated learning of unknown model terms with less data than the neural network alone. Nevertheless, estimating parameters for these hybrid models remains difficult with sparse data and limited sampling durations that are common in biological applications. In this work, we propose the use of physiology-informed regularization, penalizing biologically implausible model behavior to guide the UDE towards more physiologically plausible regions of the solution space. In a simulation study we show that physiology-informed regularization not only results in a more accurate forecasting of model behaviour, but also supports training with less data. We also applied this technique to learn a representation of the rate of glucose appearance in the glucose minimal model using meal response data measured in healthy people. In that case, the inclusion of regularization reduces variability between UDE-embedded neural networks that were trained from different initial parameter guesses.

**Author summary:** Systems biology concerns the modelling and analysis of biological processes, by viewing these as interconnected systems. Modelling is typically done either using mechanistic differential equations that are derived from experiments and known biology, or using machine learning on large biological datasets. While mathematical modelling from biological experiments can provide useful insights with limited data, building and validating these models takes a long time and often requires highly invasive measurements in humans. Efforts to combine this classical technique with machine learning have resulted in a framework termed universal differential equations, where the model equations contain a neural network to describe unknown biological interactions. While these methods have shown success in numerous fields, applications in biology are more challenging due to limited data-availability, high data sparsity. In this work, we have introduced physiology-informed regularization to overcome these instabilities and to constrain the model to biologically plausible behavior. Our results show that by using physiology-informed regularization, we can accurately predict future unseen observations in a simulated example, with much more limited data than a similar model without regularization. Additionally, we show an application of this technique on human data, applying a neural network to learn the appearance of glucose in the blood plasma after a meal.

## Introduction

The high complexity of the internal regulation of homeostasis in the human body, combined with increasing levels of detail of biological datasets, often makes direct interpretation of measurements difficult. In systems biology, this challenge is approached through dynamic mathematical modelling, which aims to systematically combine known biology and hypotheses with quantitative data. [1] An advantage of representing a biological system as a set of mathematical equations is that simulations can be performed rapidly using numerical methods providing insights into how individual interactions can give rise to systemic behavior over time and allowing for in silico hypothesis testing.

The model development cycle, whereby models are constructed, iteratively refined, and validated by comparison with experimental data, provides a practical basis for computer-assisted biology research. In this way, biological interactions which are difficult to isolate in *in vitro* experiments can be uncovered. [2, 3] For example, a recent model revealed the context-specific mechanism by which the hypoxia-inducible factor 1*α* (HIF1*α*) regulates lipid-accumulation [4] and a model of the liver X-receptor (LXR) revealed that the origin of LXR-induced hepatic steatosis lies in an increase of the free fatty-acid influx in the liver. [5] However, the model development cycle is highly laborious and time consuming, and larger dimensionalities of data complicate the construction of reliable mathematical models. [1]

With the introduction of high-throughput sequencing technology, the number of features in medical data is growing quickly. However, collecting data at the time resolution and duration necessary to train many commonly used machine learning methods is often infeasible in practice. Furthermore, while the generation of data with wearable devices achieves sufficient temporal resolution, they are still limited to very specific features, such as continuous glucose monitoring or physical activity monitoring with smartwatches. A promising solution is to integrate previously built mechanistic models, strongly rooted in known biology, with highly flexible machine learning approaches. During training, the mechanistic model simulates the known biology underlying the system, allowing the machine learning model to learn only the unknown relationship from the data. This technique is labeled Universal Approximator Differential Equation (UDE) modeling, referring to the universal approximation property [6] of neural networks. [7, 8] These universal differential equations have been shown to recover missing interactions in examples from materials science, astrophysics, and epidemiology, and have been successfully applied in environmental sciences, capturing missing terms in glacier ice flows and climate models. [9, 10]

The introduction of a neural network introduces a large number of parameters into an already complex non-linear biological model and therefore heavily influences the already complicated loss landscape of a parameterized dynamic model. When combined with limited data availability due to practical limitations of collecting data from humans, this presents itself as an increase in local minima in the loss function, having the potential to negatively impact the ability of the model to find a generalizable solution.

This challenge of training a universal differential equation system in situations where limited data is available is an active area of research. An approach by Roesch et al. proposed a two-stage approach to reduce the probability of a model getting stuck in a local minimum. [11] However, because this method initially relies on splines of the data, it is sensitive to low sampling frequencies as often encountered in biomedical data. [12] Additionally, data needs to be available for all state variables in the model, which is rarely the case in biological systems. Vortmeyer-Kley et al. instead suggested a modification to the loss function, minimizing the angle between the predicted and given points in the state-space of the system, in addition to the difference between the two points. [13]

Additionally, Turan and Jaschke proposed splitting the data into smaller parts and computing the loss of the ODE system for each part separately. This approach involves adding pseudo-initial values for all additional ODE solves into the parameter vector, and penalization of possible discontinuity between different adjacent sections. This approach is labeled multiple shooting. [14] Even though these approaches can lead to improvements in neural network behavior within differential equation systems, these solutions depend on having either a sufficient sampling frequency or a sufficient sampling duration, and do not take advantage of the wealth of knowledge available for many of these systems.

Instead, we propose to constrain the model solution to physiologically plausible regions by incorporating this additional biological knowledge in a simple fashion through physiology-informed regularization. In a general sense, regularization is a technique in machine learning that aims to reduce overfitting. In this study, physiology-informed regularization is implemented as a form of Tikhonov regularization, where the cost function is supplemented with terms penalizing biologically undesirable behaviors, such as negative metabolite concentrations or metabolites disappearing or being created out of nothing. In addition to reducing overfitting of the neural network, this form of regularization promotes the identification of biologically plausible behaviors in these larger systems. The type of biological information that is included in this way can vary per system, depending on the knowledge of the system, the available data, and the specific term that the neural network is representing. This type of physiology-informed regularization has been applied previously in classical ODE modelling [3, 15], and a similar technique was recently used in an epidemiological model using the UDE approach [16]. However, the impact of physiology-informed regularization on the training process in situations of varying data sparsity and sampling time, as well as varying the strength of different regularization terms has not been investigated previously in universal differential equation models.

In this work, we implement simple forms of physiology-informed regularization to aid in the training of two UDE systems. Firstly, we add a penalty to prevent negative concentrations in the context of a simple model describing the conversion of one molecule to another via Michaelis-Menten kinetics. Additionally, we investigate the effect of physiology-informed regularization in scenarios with varying sampling duration and scarcity. To demonstrate the benefit of physiology-informed regularization beyond a simulated example, we also applied it to a second scenario, learning the rate of appearance of glucose from a meal from human data in Bergman’s glucose minimal model [17], where the UDE term is constrained to be non-negative and have an area under the curve of glucose influx in the blood plasma that corresponds to the amount of ingested glucose.

## Materials and methods

### General structure of a UDE model

For state-variables **u**(*t*), a universal differential equation (UDE) can be formulated as:

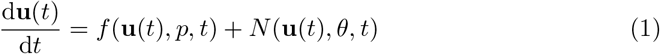

Here, *f* (**u**(*t*), *p, t*) represents the known terms in the differential equation model using the state variables **u**(*t*), over the time *t* and possibly *a priori* unknown parameters *p*.

In a typical parameter estimation procedure, these parameters *p* are estimated from data by minimizing the difference between the data and model predictions using numerical optimization techniques. For a UDE, the neural network, shown in equation 1 as *N* (**u**(*t*), *θ, t*), where *θ* indicates the neural network parameters, represents unknown terms. Similarly to the conventional parameter estimation procedure, the parameters of the UDE model are estimated from data using numerical optimization. Both *p* and *θ* can be estimated simultaneously. [18]

### Regularization in a simulated model of michaelis-menten kinetics

To test the effects of regularization on the quality of the UDE model fits to data a simulation study was performed, allowing the error of the UDE model with respect to a known ground truth to be determined.

### Model description

In this simulation, the known model consists of two ordinary differential equations (ODEs) coupled by a saturable enzymatic conversion term implemented as a Michaelis-Menten equation, which is commonly found in models of biological pathways. [19] The Michaelis-Menten model used here can be described mathematically as

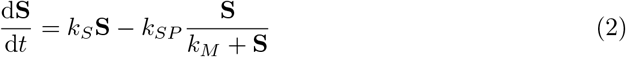

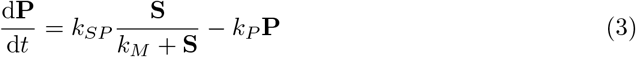

Where **S** and **P** represent the concentrations (in mM) of two distinct metabolites or molecules, **P** is produced from **S** through an enzymatic conversion with rate constant *k*_*SP*_ and Michaelis-Menten coefficient *k*_*M*_ . Furthermore, metabolite **S** stimulates its own production through rate parameter *k*_*S*_, and metabolite **P** decays linearly with its concentration and the decay rate *k*_*P*_ , displayed graphically in Fig. 1A. All parameters and initial conditions for the ground truth model simulation are described in Table 1. The time variable was taken to be measured in minutes. This and all other models in this work were implemented and simulated using Julia version 1.10 [20] and the Tsit5-solver [21] in the DifferentialEquations package. [18]

**Table 1.** Parameter values and initial state values used for data simulation of the Michaelis-Menten model.

| Parameter | Value | Unit |
| --- | --- | --- |
| $k_S$ | 0.05 | $\text{min}^{-1}$ |
| $k_P$ | 0.2 | $\text{min}^{-1}$ |
| $k_{SP}$ | 0.08 | $\text{mM} \cdot \text{min}^{-1}$ |
| $k_M$ | 1.1 | $\text{mM}$ |
| $S_0$ | 2 | $\text{mM}$ |
| $P_0$ | 0 | $\text{mM}$ |

**Fig 1.**
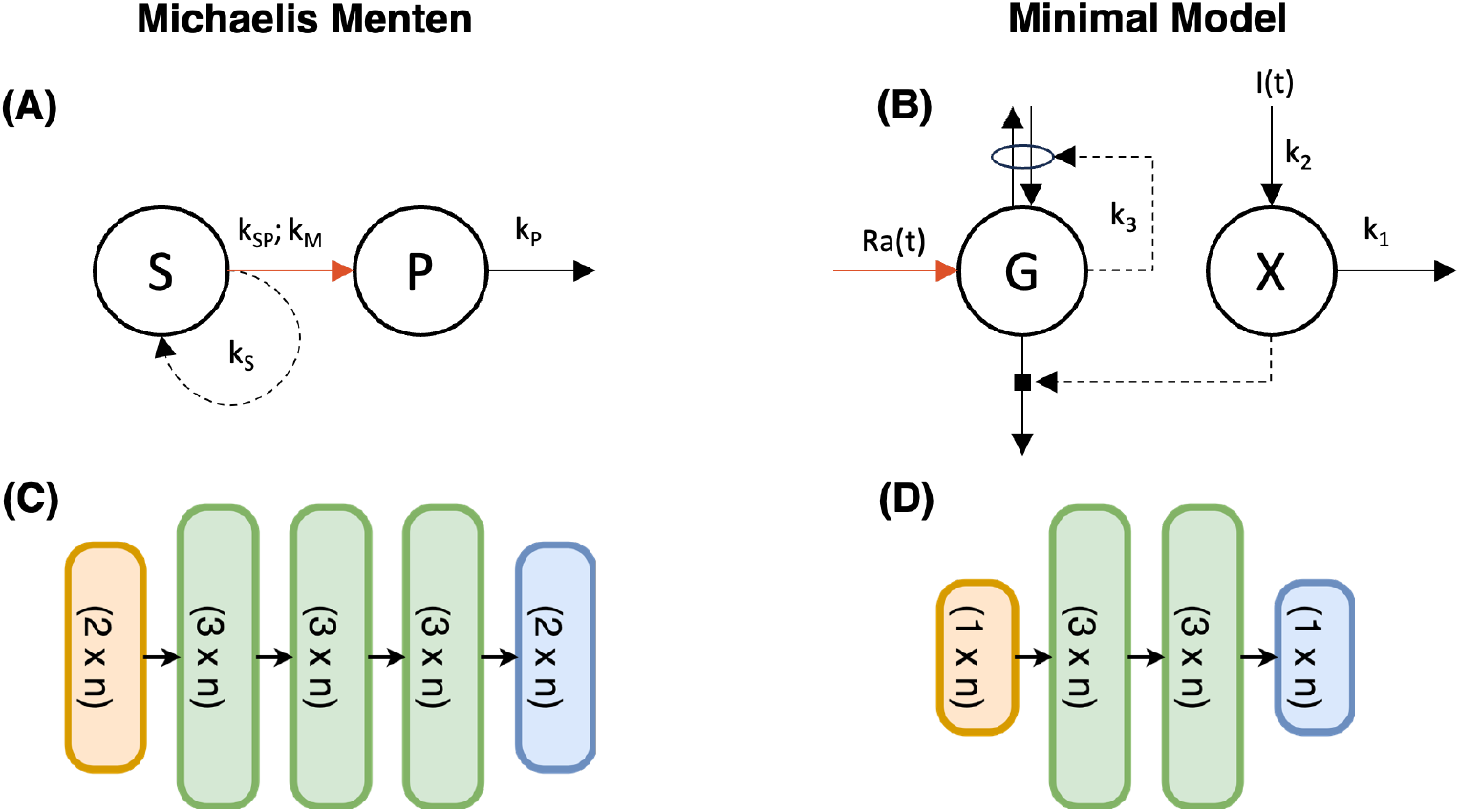
Overview of implemented model structures and architectures of incorporated networks. A: Graphical representation of the simulated model with Michaelis-Menten kinetics. B: Graphical representation of the Bergman Minimal Model. Interactions are given using dashed arrows, while fluxes are shown in solid arrows. The location of the neural network in both models is indicated by the orange arrow. C: Neural network architecture for the Michaelis-Menten model. D: Neural network architecture for the Minimal Model. All layers are densely connected.

### Data description

Data was simulated using the model and parameters described in table 1. The simulated data was split into high-resolution validation data, sampled every 0.1 time units between 0 and 400 time units, and several lower resolution training datasets. Each training dataset represents a combination of values set for the sampling step size and the duration of sampling. Samples were extracted at intervals of 5, 10, or 20 minutes for sampling durations of 20 to 100 minutes. Additionally, noise was added to each training dataset to simulate measurement conditions. The noise distribution was assumed to be normal, with a standard deviation of 5% of the maximum value in the validation dataset of the corresponding state-variable. In the cases where addition of noise resulted in negative measurements, measurement values were set to 0.

### Construction and training of the UDE model

The UDE model is formed by removing the michaelis-menten coupling term and replacing it with a single fully connected neural network with two inputs, three hidden layers of size 3 with Gaussian radial basis activation functions, and a single output with linear activation (Fig. 1C). The gaussian radial basis activation function can be formulated as

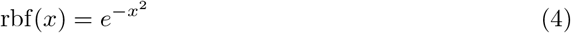

In this case, the parameter values for *k*_*S*_ and *k*_*P*_ were assumed to be known and fixed to the values in Table 1. For implementation of the neural network, the Lux.jl package [22] was used.

Using this implementation, a loss function for estimating the neural network parameters was formulated as

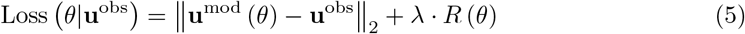

Indicating the sum of squared differences between model predictions (**u**^mod^ (*θ*)) for neural network weights *θ* and data (**u**^obs^), combined with a regularization function *R* (*θ*), of which its contribution is controlled by a scaling hyperparameter *λ*. As such, the case where *λ* = 0, represents an unregularized setting. The regularization function was chosen to prevent the neural network from predicting negative concentrations of either state variables **A** or **B**, resulting in a one-sided penalty term formulated as

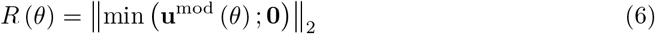

In this formulation, any positive value for the model predicted concentration remains unpenalized, while the penalty on negative concentrations grows linearly with the square of their magnitudes.

Estimation of neural network parameters was achieved by performing gradient-based optimization, starting from a random initial set of parameters, to find a parameter set that minimizes the loss function. To improve convergence, a two-stage optimization procedure was used, starting out with the Adam [23] optimizer with a learning rate of 0.01 for 500 iterations, followed by the Broyden-Fletcher-Goldfarb-Shanno (BFGS) [24–27] algorithm in the second stage. The BFGS algorithm was used with an initial step norm of 0.01, running until convergence based on a function tolerance of 1e-6 and an input tolerance of 1e-6, or until the maximum number of iterations, set at 1000, was reached. The optimization procedure was carried out in Julia version 1.10 using the Optimization package. [28]

For each simulated training dataset, optimization was performed 100 times using random initialization of the neural network weights. This procedure was carried out using regularization strengths of 0 (no regularization), and 10^*x*^ for *x* ∈ {−9, −8, −7, −6, −5, −4, −3, −2, −1}.

### Evaluation of regularization in simulated conditions

To investigate the effect of the regularization penalty, model performance was evaluated by computing an evaluation error based on the simulated validation dataset as

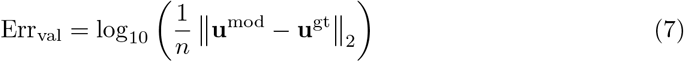

The value of n represents the number of validation points, which was set to a value of 4001, corresponding to the amount of points in the generated validation dataset.

### Effect of physiology-based regularisation on human data

In addition to the simulation study, the effect of physiology-based regularization on the training of a UDE model was also evaluated in a real-life setting. As a test case, a neural network was introduced into the oral glucose minimal model [17, 29] to learn a representation of the rate of glucose appearance from the gut and was trained using a large public dataset of meal responses.

### Model description

The glucose minimal model describes the dynamics of plasma glucose regulation by insulin at a course-grained level and is formulated as

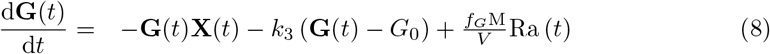

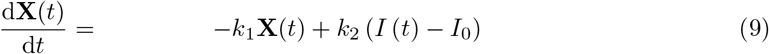

Where **G** and **X** describe plasma glucose concentration and a generic glucose lowering effect respectively. The model has two forcing functions, Ra (*t*) describing the appearance of glucose from a meal in the plasma, and *I* (*t*) for incorporating the measured plasma insulin values. *G*_0_ and *I*_0_ denote the fasting glucose and insulin concentrations respectively, which establish the set-points of the model. *M* represents the mass of glucose in the meal in milligrams (mg), *f*_*G*_ is a conversion factor from mg to mmol glucose, and *V* is the distribution volume of glucose in liters, computed from the body weight as in [15] using

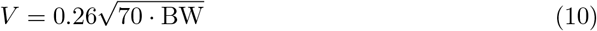

Where BW refers to the body weight. Used values for *M* , *f*_*G*_, and *V* are included in table 2. The model has three kinetic parameters, *k*_1_, *k*_2_, and *k*_3_ which describe the rate of decay of **X**, the production of **X** as a function of the plasma insulin measurement, and the rate of a general insulin-independent plasma glucose removal, respectively. A visual representation of the model is shown in Fig. 1B.

**Table 2.** Parameter and constant values used for the glucose minimal model.

| Variable | Value | Unit | Description |
| --- | --- | --- | --- |
| $f_G$ | $5.551 \cdot 10^{-3}$ | $\text{mmol} \cdot \text{mg}^{-1}$ | Conversion of glucose from mg to mmol |
| $M$ | $85.5 \cdot 10^3$ | mg | Mass of glucose in the meal in mg |
| $V$ | 18.57 | L | Volume of distribution of glucose in the blood plasma |
| $k_1$ | $4.91 \cdot 10^{-2}$ | $\text{min}^{-1}$ | Rate of decay of insulin action |
| $k_2$ | $2.75 \cdot 10^{-5}$ | $\text{mL} \cdot \mu\text{IU}^{-1} \cdot \text{min}^{-1}$ | Production of insulin action based on plasma insulin |
| $k_3$ | $4.61 \cdot 10^{-2}$ | $\text{min}^{-1}$ | Rate of insulin independent glucose removal |

### Data description

To train this model the meal response data for 1,002 individuals from the PREDICT UK study were acquired from the paper by Berry et al. [30] Population demographics are shown in Table 3. The data contained plasma glucose and insulin concentrations measured at 0, 15, 30, 60, 120, 180, and 240 minutes following the consumption of a standardized solid meal. The meal contained 85.5 g of carbohydrates, 52.7 g of fat and 16.1 g of protein. Individuals with missing values for glucose or insulin in any time point for this meal were removed. Data from the remaining 903 subjects is averaged across individuals for each time point. This average meal response data is used to train the UDE-glucose model.

**Table 3.** Population demographics of the PREDICT UK cohort.

| Variable (unit) | Value |
| --- | --- |
| Sex (M/F) | 279/729 |
| Age (Y) | 45.48 ± 11.88 |
| Body weight (kg) | 72.88 ± 15.27 |
| Height (cm) | 168.53 ± 10.51 |
| BMI (kg m<sup>-2</sup>) | 25.59 ± 5.02 |
| Fasting plasma glucose (mM) | 4.96 ± 0.48 |
| Fasting plasma insulin (mU L<sup>-1</sup>) | 6.13 ± 4.27 |

Where applicable, values are shown as mean ± standard deviation. Values adapted from Berry et al. [30]

### Construction and training of the UDE model

In this instance, the UDE approach is deployed to estimate the plasma glucose appearance (Ra (*t*)) in the glucose-minimal model. Only the rate-of-appearance function is estimated from data, while values for the model parameters *k*_1_, *k*_2_, and *k*_3_ are assumed to be known. To obtain values for the parameters *k*_1_, *k*_2_, and *k*_3_, an initial rate-of-appearance function was assumed using a mechanism described by Korsbo et al. [31] (see S1 Rate of Appearance). *k*_1_, *k*_2_, *k*_3_ were estimated through least-squares estimation on the PREDICT data and subsequently fixed to the values depicted in table 2.

The neural network used to estimate Ra was a fully connected network with one input, two hidden layers of size 3 with Gaussian rbf activations (equation 4) and one output layer with linear activation (Fig. 1D). In addition to the nonnegativity regularization, as used in the simulated Michealis-Menten condition, an additional regularization penalty was incorporated to the training of the glucose-minimal UDE to ensure the area under the rate-of-appearance curve should equal 1. This penalty ensures that all the glucose from the provided meal would be taken up into the plasma compartment of the model.

The values for regularization strengths were set to either 0 (no regularization) or a value in 10^*x*^ with *x* ∈ {−2,−1, 0, 1, 2} for both types of regularization. Optimization was performed for all possible combinations of both the nonnegativity regularization and the area-under-curve regularization. The neural network parameters were initialized with random parameters sampled from a zero mean gaussian distribution with a standard deviation of 1 · 10^−4^. For each experimental condition, 100 initializations were optimized.

## Results

### Regularization in a simulated model of michaelis-menten kinetics

Michealis-Menten UDE models were trained for different regularization strengths, and sampling schedules. Fig. 2 visualizes 25 model fits with the lowest training error for models trained both with (*λ* = 10^−5^ and *λ* = 1) and without regularization for a selected sampling duration of 40 minutes with samples taken every 5 minutes. The grey area indicates where data has been supplied for parameter estimation. Comparing the model fits to the ground truth model (black, solid and dashed), we see that in the regularized cases (B) and (C) there is good correspondence between the model simulation and the ground truth, while the unregularized models (A) fail to predict the concentrations correctly beyond the supplied data. Furthermore, the interquartile ranges are narrower for the models trained with regularization indicating a lower variance in model prediction. Differences between regularization strengths are also observed. The correspondence between model and simulated true values is higher for *λ* = 10^−5^ than for *λ* = 1.

**Fig 2.**
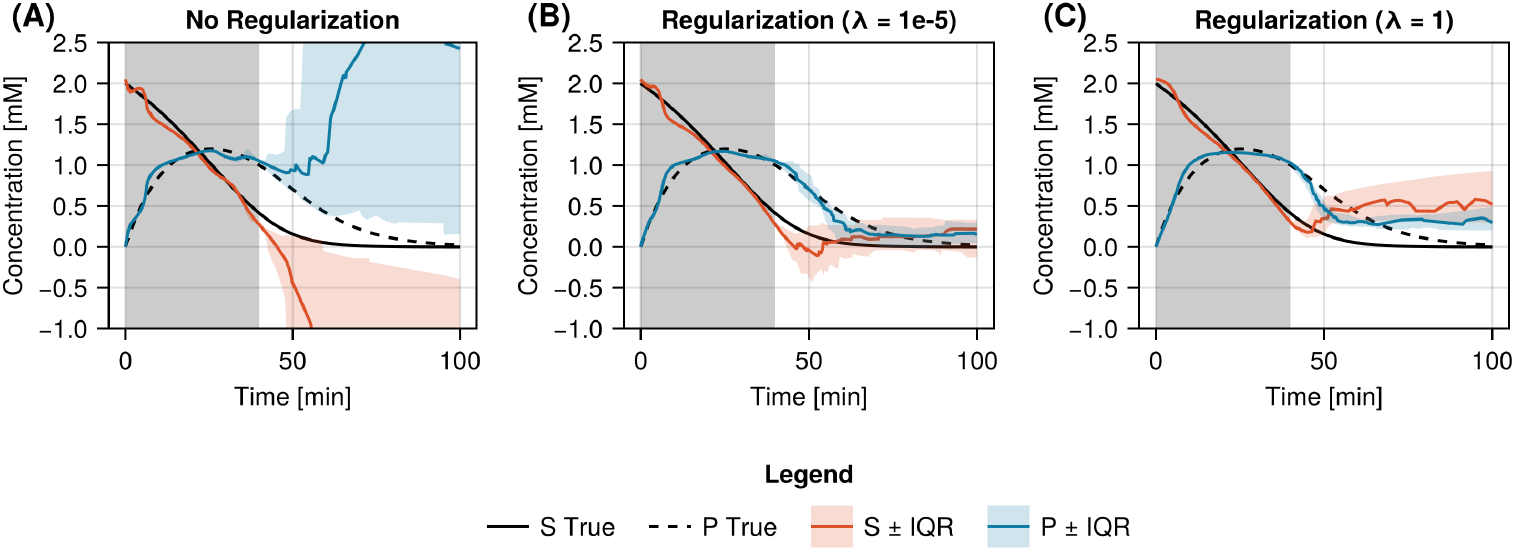
Visualization of learned Michealis-Menten models trained with and without regularization. Median predictions for the Michealis-Menten model for species S (red) and species P (blue) including the first and third quartiles for A: the case without regularization (*λ* = 0), B: mild regularization (*λ* = 10^−5^), and C: strong regularization (*λ* = 1) for a sampling duration of 40 minutes. The red and blue shaded regions indicated the interquartile range of the top 25 models, selected based on the training error, for species S and P respectively.. The sampling duration used for training is marked in the grey shaded region. The ground truth model is also visualized with the black solid (S) and dashed (P) lines to allow comparison of the model fits.

Additionally, we investigated the distributions of the ground-truth evaluation error (Err_val_) for these initializations, which are shown in Fig. 3. In this case, the used training set was sampled every five minutes for 40 units of time. The figure shows a clear bimodal distribution when the models are trained without regularization, with the majority of model fits finding an erroneous steady state with a large Err_val_, whereas for all regularization strengths the majority of the model initializations have a low validation error, as evident with the peak close to an E_val_ of 4. The distributions of all regularized cases are similar in location, but differ slightly in shape. For stronger regularization strengths, the peak around Err_val_ = 4 becomes sharper.

**Fig 3.**
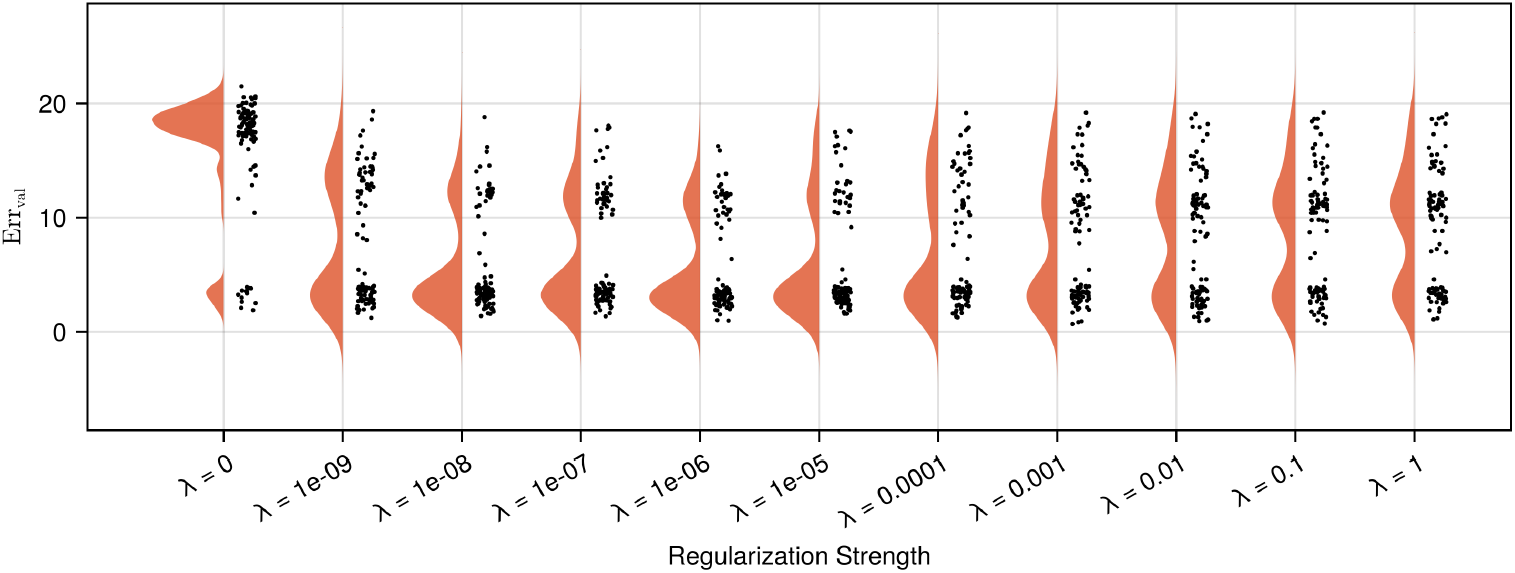
Distributions of the validation error for Michealis-Menten models trained with different regularization strengths. The distributions for of validations errors (E_val_) computed on each of the 100 models for different regularization strengths. The models are trained using data over a sampling duration of 40 units of time ,The validation errors (E_val_) are computed on the simulated ground-truth data over a duration of 400 units of time. The validation error is presented on a log10 scale. For models trained without regularization (*λ* = 0) a biphasic distribution in validation errors is found, with the majority of model fits finding a local minima with a large Err_val_. The incorporation of regularization (*λ* ≥ 0.1) during the training of the Michealis-Menten UDE improves the long-term dynamics of the learned model terms with the majority of validation errors located in the vicinity of 4 Err_val_. The effect of the regularization strength on the performance is mainly seen in the size of the peak around Err_val_ = 4, which becomes larger for stronger until *λ* = 10^−5^ and then flattens out, as more model fits remain at larger validation error values.

We also evaluated the effect of data sampling duration and sampling sparsity on the quality of the learned model. Fig. 4 shows the mean evaluation errors computed for the unregularized case (orange) and two regularized cases (*λ* = 10^−5^ and *λ* = 1) (blue and green respectively). The sampling duration varied from 20 to 100 minutes in steps of 10 minutes for samples taken every 5 and 10 minutes. For samples taken every 20 minutes, the sampling duration varied from 40 to 100 minutes in steps of 20 minutes.

**Fig 4.**
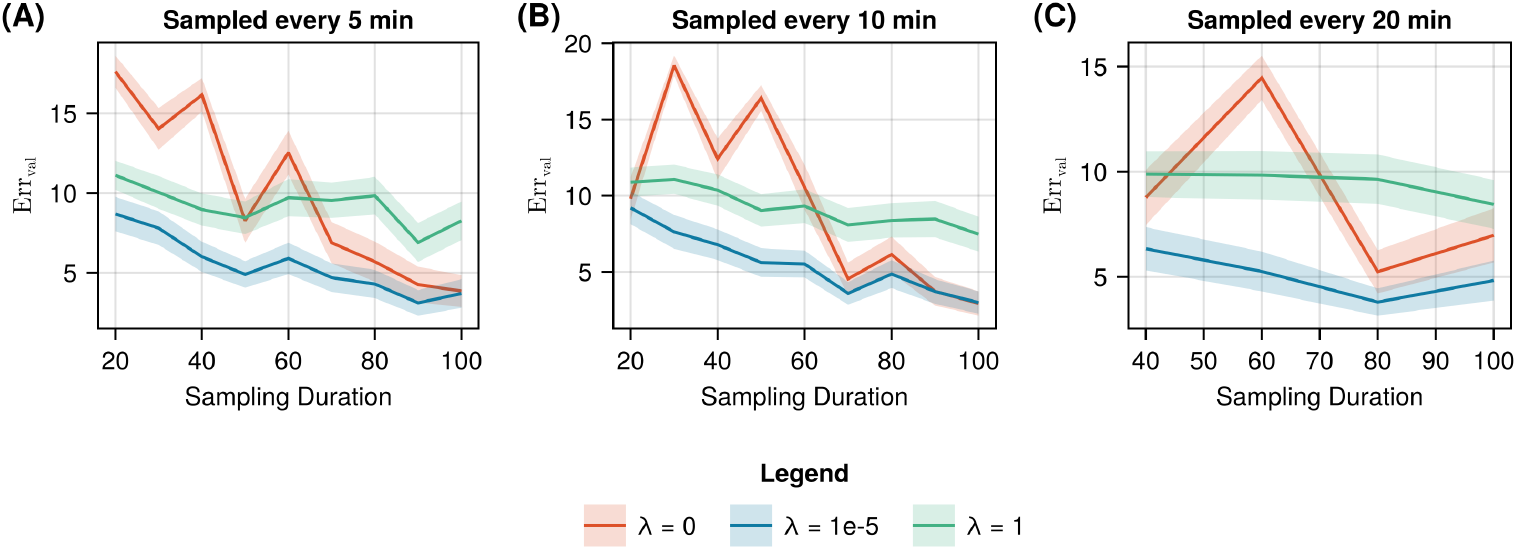
Mean validation error for varying durations of data collection and sampling frequencies with and without regularization. Mean (solid line) and standard deviation (shaded) of the validation error of the optimized models with slight (*λ* = 10^−5^), strong (*λ* = 1), and without regularization, for different sampling durations of the training data sampled every A: 5 minutes, B: 10 minutes, and C: 20 minutes.

The values in Fig. 4A show that the average model reconstruction error in the regularized case does not seem strongly influenced by the sampling sparsity, whereas the increase in sampling duration reduces the validation error. For slight regularization, we observe the lowest validation error, consistently outperforming the average unregularized models up until a sampling duration of 80 minutes, after which the average performance of both conditions remains similar. For strong regularization, we observe a lower validation error than the unregularized case before 60 minutes of sampling duration, after which the unregularized case shows lower validation errors.

### Effect of physiology-based regularisation on human data

To evaluate the generalizability of the benefit of physiology-based regularization on the training of UDE models, we also applied the approach in a second example using real data. In this instance to enable simulation of the glucose response to a complex meal we aimed to use a universal approximator differential equation to learn an adjusted glucose rate-of-appearance in the glucose oral minimal model.

Hyperparameter values for both nonnegativity and area-under-curve regularization were evaluated (S2 Fig). The fits of the top 25 models based on training error for models train with (*λ*_nonneg_ = 100 and *λ*_AUC_ = 1) and without regularization are shown in Fig. 5A and B. In both conditions the models can be seen to fit the average data. Moreover, following the meal, the models trained with regularization tend to converge to a steady-state approximately equal to the fasting plasma glucose concentration measured prior to the meal. Whereas the models trained without regularization either find no steady-state or a steady-state over a range of 1 mmol/l around the fasting plasma glucose. In Fig. 5C and D, the rate-of-appearance curves for each of the models are shown. Both curves show a biphasic glucose appearance. At the start of both curves, the regularized curve is consistently positive, while some unregularized curves become negative. Furthermore, the observed difference in steady-state in the plasma glucose curves between the regularized and unregularized cases can also be seen in the rate-of-appearance curves. In Fig. 5E and F, the standard deviation and interquartile ranges of the rate-of-appearance curves are visualized over time, demonstrating that physiology-based regularization improves long term stability of the learned solution as the steady state standard deviation and interquartile range remain low. In the unregularized case, both metrics increase after the final glucose data point.

**Fig 5.**
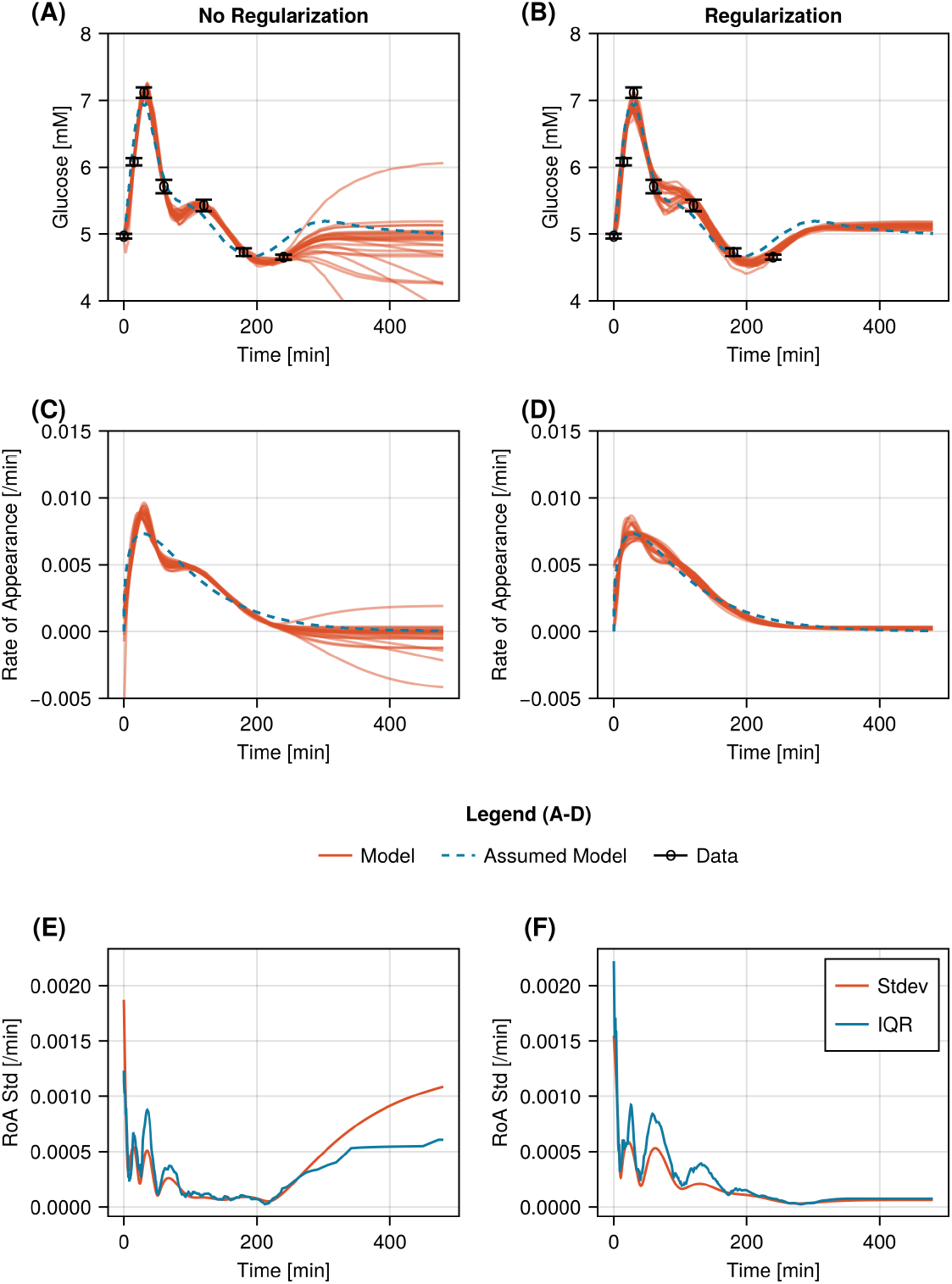
Results of the 25 glucose-minimal UDEs with the lowest training error values for models trained with and without regularization. A: Fits of models trained without regularization (red) and the original rate-of-appearace (blue dashed) to measured plasma glucose concentrations, where data is shown as mean (black circles) ± standard error. B: Fits of models trained with regularization (red) and the original rate-of-appearace (blue dashed) to measured plasma glucose concentrations, where data is shown as mean (black circles) ± standard error. C: Learned rate-of-appearance functions for the 25 glucose-minimal UDEs with the lowest training error values trained without regularization (red) and the original rate-of-appearance function (blue dashed). D: Learned rate-of-appearance functions for the 25 glucose-minimal UDEs with the lowest training error values trained with regularization (red) and the original rate-of-appearance function (blue dashed). E: Standard deviation (red) and interquartile range (blue) of the rate-of-appearance function over time for the 25 unregularized models with the lowest training error values. F: Standard deviation (red) and interquartile range (blue) of the rate-of-appearance function over time for the 25 regularized models with the lowest training error values.

To verify that regularization on the area under the curve results in conservation of glucose mass, the AUC of the learned rate-of-appearance curves was computed for models trained both with and without regularization. Fig. 6 shows that the regularization strongly forces the AUC of the rate of glucose appearance to 1.0, with some models showing an AUC between 0.9 and 1.0, while the spread of the unregularized AUC values is markedly larger.

**Fig 6.**
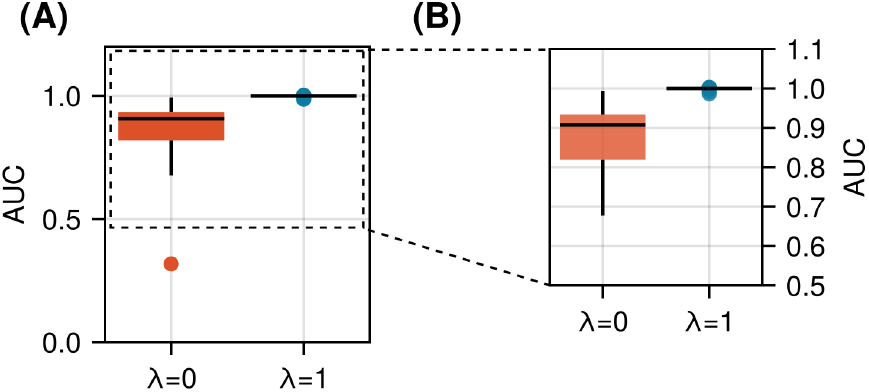
Area under the rate-of-appearance curve for the regularized and unregularized models. A: Distribution of the area under the curve of the unregularized (red) and regularized (blue) models. B: Detailed view of the top part of the figure in A.

All code and data to reproduce the results are available from the GitHub repository linked to this publication (https://github.com/Computational-Biology-TUe/ude-regularization).

## Discussion

While methods combining machine learning with knowledge-based models, such as UDEs, have great potential to rapidly advance modelling in systems biology, the increased model complexity coupled with the number of trainable parameters still limits their use to either well-studied systems, or cases where abundant data is readily available. In this study, we demonstrate the benefit of physiology-informed regularization as a technique to both stabilize training of UDEs for biological systems, as well as increase the biological plausibility of the found solutions.

We have demonstrated the advantage of regularization techniques in a simulated example, where the ground truth was available. In the simulated case, we show that regularization supports successful training of a UDE model in low data scenarios. Furthermore, distributions of validation errors from different initializations indicated that regularization increases the likelihood of learning more accurate models. Additionally, we have demonstrated that the effects of physiology-informed regularization are strongest with shorter sampling durations, reflecting the ability of the regularized model to fit the system with fewer information. However, we also show the necessity for hyperparameter tuning, as a too strong regularization can negatively impact the forecasting and generalisation abilities of the model. We demonstrate that incorporating physiology-informed regularization shows benefit when applied on real-world data, by applying it to learn a representation of the rate of glucose appearance from a meal in the glucose-minimal model using meal responses measured in healthy humans.

However, in reality the ground truth is often unavailable. Therefore, multiple initial parameter sets are often used to increase the reliability of learned models. In this study, we trained 100 models for each scenario using random parameter initialization. We used the training error to construct a population of top performing models. When using regularization during model training, the population’s capacity to predict behaviors beyond the training data consistently outperformed the population of models trained without regularization (Fig 2, Fig 5 A and B). Furthermore, in both simulated and real-life situation regularization was shown to reduce the variability in found solutions with the lowest training error. In the real-life situation, this effect is smaller than in the simulated case. Furthermore, based on the differences between the standard deviation and the interquartile range in the real-life situation it is likely that the continued increase in standard deviation is driven mostly by outlier model fits, while the eventual stabilization of the increased interquartile range reflects the greater distribution in learned steady-state values.

This work extends earlier applications of physiology-informed regularization [3, 15] into the domain of scientific machine learning, and provides a simple alternative to earlier methods to improve UDE stability during training. [11, 13, 14] As basic physiological or chemical constraints are often available in biological models, the extension of a UDE training process with physiology-informed regularization is often straightforward. While previous methods of improving UDE training have shown their efficacy in other engineering fields, the limited data availability often encountered in biological applications makes the application of these methods challenging. By accounting for straightforward biological constraints, the physiology-based regularization approach presented in this work addresses the training stability by providing additional information about the system, as well as guiding the gradient of the cost function towards biologically more plausible solutions.

Another method that bears resemblance to the type of regularization demonstrated here, is ridge regression, which instead of constraining model behavior, pulls parameters towards predetermined values during model training and is often used in purely data-driven machine learning applications to limit model overfitting on uninformative features by penalizing the size of the parameters directly. However, in context of neural networks, the individual parameters have limited meaning by themselves. While a form of ridge regression can still be useful in UDE training to prevent strong overfitting on the data, this does not prevent the model from learning biologically implausible behavior. Nevertheless, if necessary, both ridge regression and physiology-informed regularization could also be used in conjunction, noting of course that this would introduce an additional hyperparameter to be tuned.

While the results indicate clear benefits of regularization for training of UDE models for biological systems, all model fits in the real-life scenario were performed on the average glycemic response. We have not tested regularization in a setting where models were fit to individual meal responses, as the stronger contribution of measurement noise to that type of data complicates solid assessment of regularization techniques. Furthermore, we have fixed the parameters of the meal response model to the values estimated using the hypothetical rate-of-appearance function. As a result, we may bias the learned neural rate-of-appearance towards a specific shape generated by the initial glucose-minimal model. However, this limitation is present in both regularized and unregularized models trained here. In the future, this could potentially be investigated in further research in context of the capabilities of neural networks within UDE models to correct for errors in model parameters outside of the neural network. Furthermore, while the inclusion of this neural network into the system provides satisfactory forecasting abilities, the interpretability of the model is reduced. To improve interpretability, tools like symbolic regression [32, 33] could be applied after training, to derive a functional representation of the missing model component. [7]

Additionally, while we have demonstrated the benefits of physiology-informed regularization in both a simulated and real data setting, the models evaluated in this work are relatively simple, describing interactions between two state variables. In future work, the efficacy of physiology-informed regularization should be confirmed in larger model systems.

In summary, we have presented physiology-informed regularization as a simple, yet powerful and generalizable approach for the improvement UDE training in biological models. The used of physiology-informed regularization not only improves long-term predictive stability, reducing model variance. But it can also be seen to reduce non-physiological behavior in the neural network component.

## Acknowledgements

The research presented in this manuscript was supported by a Starting Package from the Eindhoven AI Systems Institute (EAISI) awarded to S.O’D. N.A.W.v.R is supported by a grant from the Dutch Research Council (NWO) [https://www.nwo.nl/] as part of the Complexity Program (project number 645.001.003).

## Supporting information

**S1 Rate of Appearance** Given the absence of model equations to describe the rate of glucose appearance from the meal (Ra (*t*)), a delayed pulse input function was used to describe plasma glucose appearance for parameter estimation. This function is derived from a glucose pulse input function (dirac delta) subject to a linear chain of delays with a real-valued depth *σ* and a rate parameter *k*_meal_, which is assumed to be constant across each delay compartment. Assuming that the initial meal input can be modelled using a Dirac delta, the resulting delayed glucose input is formulated as [31]

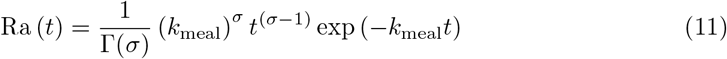

Where Γ(·) denotes the gamma function. For *σ* and *k*_meal_, the values 1.4 and 0.014 min^-1^ were used, respsectively.

**S2 Fig.**
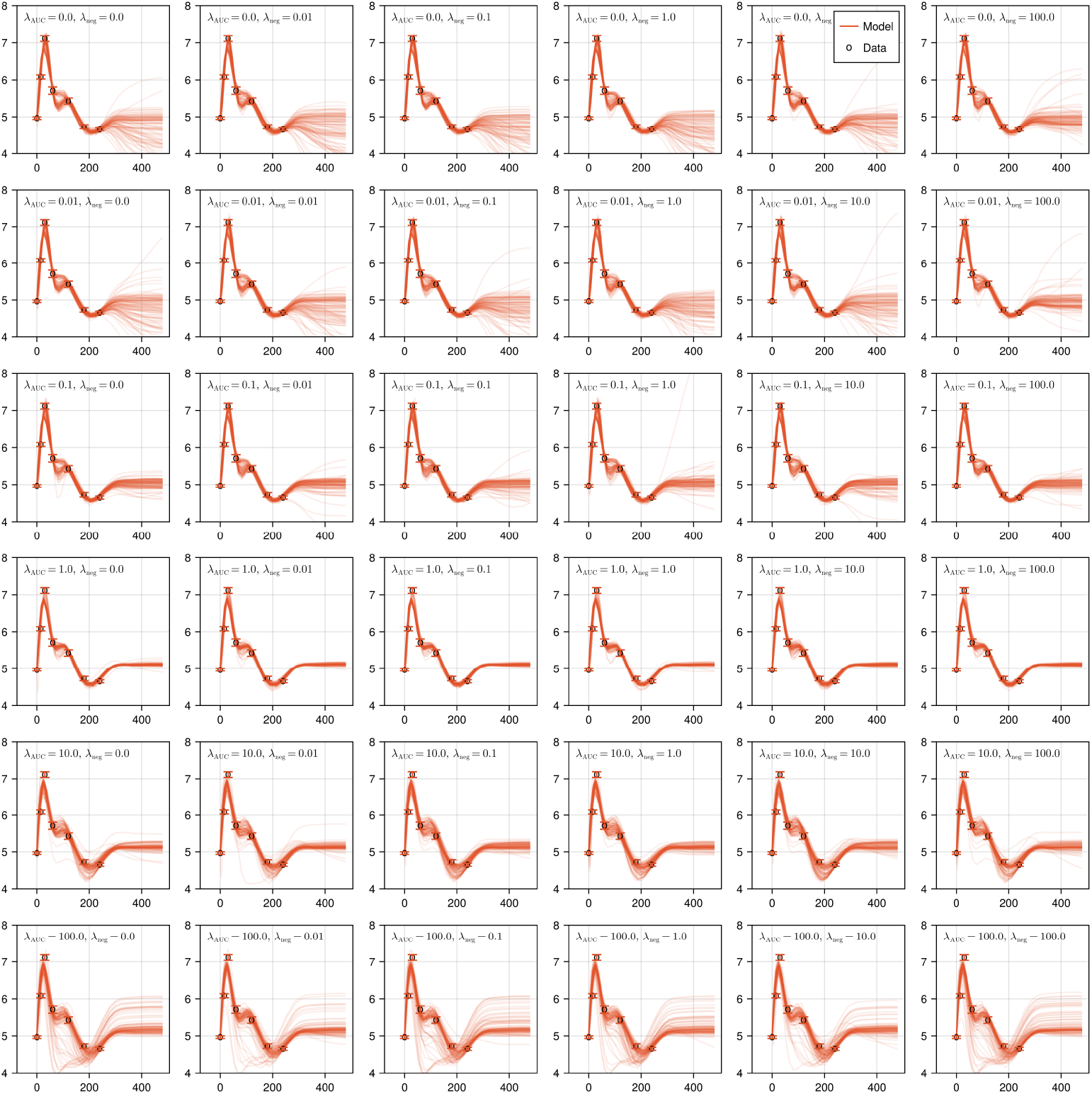
Model fits of the glucose minimal model including the neural network rate-of-appearance term for varying regularization strengths. Each solid line represents a model with a unique initial parameter set. The data is shown in circles. Each column represents the non-negativity regularization strength, while each row represents the area-under-curve regularization strength.

